# Re-Discovery of the *Phytophthora* PAMP Pep-13 Receptor Using Potato Inbred Lines

**DOI:** 10.64898/2026.03.15.709221

**Authors:** Xuefeng Fan, Dongyue Li, Lin Cheng, Yuqi Zhu, Yue Han, Tongjun Sun

## Abstract

Plants employ cell surface receptors to recognize pathogen-associated molecular patterns (PAMPs) and activate pattern-triggered immunity, a crucial defense mechanism against invading pathogens. Pep-13 is a PAMP derived from a class of conserved cell wall transglutaminases present in *Phytophthora* species, and its receptor PERU (Pep-13 receptor unit), was reported recently. In our parallel study, we observed distinct responses to Pep-13 between two diploid potato inbred lines: E4-63 recognizes Pep-13, whereas A6-10 does not. Genetic analysis demonstrated that Pep-13 recognition in E4-63 is controlled by a single genetic locus, tentatively designated *TGER* (*Transglutaminase elicitor response*). Through bulked segregant analysis sequencing, followed by complementation assays, we found that the *TGERa* gene in E4-63 is essential for Pep-13 recognition. Sequence alignment revealed that TGERa shares 99.91% amino acid sequence identity with PERU, indicating that *TGERa* and *PERU* are allelic variants of the same gene (*PERU*/*TGERa*). *TGERb*, a highly homologous gene of *TGERa*, was identified in the E4-63 genome; notably, TGERa, but not TGERb, can recognize Pep-13. We further demonstrated that TGERb exhibits defects in ligand binding. Additionally, we found that the *TGERa* allele in A6-10 is a weak allele with reduced expression levels, presumably resulting from a 3 kb DNA fragment insertion in its first intron. Heterologous introduction of *TGERa* into *Nicotiana benthamiana* and tomato significantly enhanced their resistance to *Phytophthora infestans*. Collectively, our findings confirm that PERU/TGERa functions as the Pep-13 receptor in potato and provide a valuable molecular target for improving *Phytophthora* resistance in plants.

---

Potato (*Solanum tuberosum* L.) is a major staple food crop for about 1.3 billion people worldwide (Stokstad, 2019). Its asexual propagation and complex tetraploid genome lead to long breeding cycles. To overcome this issue, a diploid potato hybrid breeding system was developed using inbred lines (Zhang et al., 2021), which accelerates breeding and provides valuable materials and genomic resources for functional genomic research in potato.

Plants mount pattern-triggered immunity (PTI) upon recognition of pathogen-associated molecular patterns (PAMPs) by cell surface receptors. Pep-13, a PAMP derived from conserved *Phytophthora* cell wall transglutaminases, is detected by the leucine-rich repeat (LRR)-receptor kinase PERU (Pep-13 receptor unit) in potato DM (Torres Ascurra et al., 2023). However, this report indicated that there are no PERU homologs in potato inbred lines A6-26 and E4-63 (Torres Ascurra et al., 2023, Supplemental data_s4). In our parallel study, we characterized Pep-13 responsiveness in the hybrid potato parental lines A6-10, A6-26, and E4-63 (Zhang et al., 2021). While Pep-13 triggered hypersensitive cell death (HR) in A6-26 and E4-63, it failed to induce HR in A6-10 (Supplemental Figure 1A), an unexpected observation given that A6-10 and A6-26 are homozygous inbreds derived from the common ancestor PG6359 (Zhang et al., 2021). Additionally, F_1_ plants of a cross between A6-10 and E4-63 responded to Pep-13, and F_2_ populations exhibited a 3:1 segregation ratio of responsive to non-responsive plants (Supplemental Figure 1B-C). These data indicate that Pep-13 recognition in E4-63 is governed by a single dominant locus, tentatively named *TGER* (*Transglutaminase elicitor response*).

Here, we identified that *TGERa* in E4-63 is essential for Pep-13 recognition. Sequence alignment revealed that TGERa shares 99.91% amino acid identity with PERU, indicating that *TGERa* and *PERU* are allelic variants of the same gene (hereafter referred to as *TGERa*). We verified that TGERa associates with immune SERKs to sense Pep-13, whereas its close paralog TGERb cannot mediate Pep-13 recognition. We also found that the *TGERa* allele in A6-10 represents a weak allele with lower expression. Finally, ectopic expression of *TGERa* markedly enhanced plant resistance to *Phytophthora* pathogens, supporting its potential as a promising molecular target for improving *Phytophthora* resistance in crops.

To identify the Pep-13 receptor in E4-63, we employed BSA-seq and transgenic complementation assays, demonstrating that *TGERa* in E4-63 is essential for sensing Pep-13 and Pep-25, a 25-amino-acid peptide containing the Pep-13 with a stronger cell-death-inducing activity (Supplemental Figures 1-3; see details in Supplemental Note 1). TGERa is a typical LRR-RK; immunoprecipitation - mass spectrometry (IP-MS) analysis in *N. benthamiana* (*Nb*) identified six SERK members, including NbSERK3a/b, NbBAK1a/b and NbSERK2a/b, as Pep-13-induced interacting partners of TGERa (Supplemental Figure 4A). Phylogenetic analysis showed that these NbSERK proteins, together with their homologs in potato (StSERK2, StSERK3A/B) and *Arabidopsis* SERKs (Ma et al., 2016), form a single clade (Supplemental Figure 4B). We further verified that Pep-13 induces interactions between TGERa and these StSERKs (Supplemental Figure 5A-C); moreover, silencing these six *NbSERKs* abrogated Pep-25/TGERa-induced cell death (Supplemental Figure 5D-J). Notably, TGERa was shown to bind biotin-labeled Pep-13 (Supplemental Figure 5K). Collectively, these data demonstrate that TGERa complexes with immune SERKs to perceive Pep-13.

Next, we compared the coding sequences (CDS) of *TGERa* and *PERU*. The TGERa^E4-63^ CDS is 78 bp longer than that of PERU^DM^, corresponding to an in-frame upstream open reading frame (uORF1) (Supplemental Figure 6A). Consequently, the annotated TGERa^E4-63^ carries an extra 26 amino acids at the N-terminus, with the rest of the sequence nearly identical except for one substitution (A317 in PERU vs. G317 in TGERa-dN26), suggesting TGERa^E4-63^ and PERU^DM^ are allelic variants of the same gene. To map the translational start site, we mutated the ATG codons of the uORF and the main ORF separately. Mutation of the main ORF ATG drastically decreased TGERa^E4-63^ protein accumulation under both the 35S and native promoters (Supplemental Figure 6B-E), indicating TGERa^E4-63^ translation mainly initiates from the main ATG. As a single-pass transmembrane protein, the N-terminal 26-amino-acid extension likely impairs signal peptide function, leading to lower protein abundance.

By contrast, the annotated PERU protein from LPH60-5 (PERU^LPH^) contains an extra 24-amino-acid insertion relative to TGERa/PERU (Supplemental Figure 7A). Although PERU^LPH^ was reported to recognize diverse Pep-25 variants, including the nearly inactive Pep-25W2A (Torres Ascurra et al., 2023), we found this insertion corresponds to the in-frame intron 2 (Supplemental Figure 7B). Intron 2 retention was detected in a fraction of *TGERa* transcripts from A6-26 leaves under both mock and *Phytophthora*-inoculated conditions (Supplemental Figure 7C-D). We then generated TGERa-In2R, which carries synonymous mutations locking intron 2 in the mature transcript (Supplemental Figure 7E-F). Unlike TGERa, TGERa-In2R failed to activate MAPK or induce cell death upon Pep-13 treatment and could not bind Pep-13 or Pep-25 (Supplemental Figure 8), indicating this intron-2-retained isoform produces a non-functional protein. Thus, the extra peptide in PERU^LPH^ likely stems from annotation errors and does not contribute to its broad ligand recognition.

Sequence alignment between the gene in construct #9 (Supplemental Figure 2) and *E454_03G001350*.*1* revealed multiple SNPs (Supplemental Table 3). A sequence identical to *E454_03G001350*.*1* was subsequently re-isolated from the amplification products of *E454* cDNA library using the same primers for *E454_03G001350*.*1* and designated *TGERb*. TGERa and TGERb share 87.9% amino acid identity, with 85.3% identity between their extracellular domains and 96.8% identity between their cytoplasmic kinase domains. Transient expression of *TGERa*, but not *TGERb*, in *Nb* leaves triggered MAPK activation and cell death following Pep-13 treatment (Supplemental Figure 9A-B). Consistently, transgenic *Nb* lines expressing *TGERa*, but not *TGERb*, exhibited cell death in response to Pep-25 (Figure 1A). Moreover, *np-TGERb* lines failed to trigger a ROS burst or activate MAPK signaling following Pep-13 treatment (Figure 1B-C; Supplemental Figure 9C). Co-IP assays further revealed that Pep-13 promoted the association of StSERK3A with TGERa, but not TGERb (Figure 1D). Unlike TGERa, TGERb did not bind Pep-13 or Pep-25 (Figure 1E-F). Together, these results illustrate that TGERb cannot perceive Pep-13/25 owing to the impaired ligand-binding activity.

**Figure 1.**
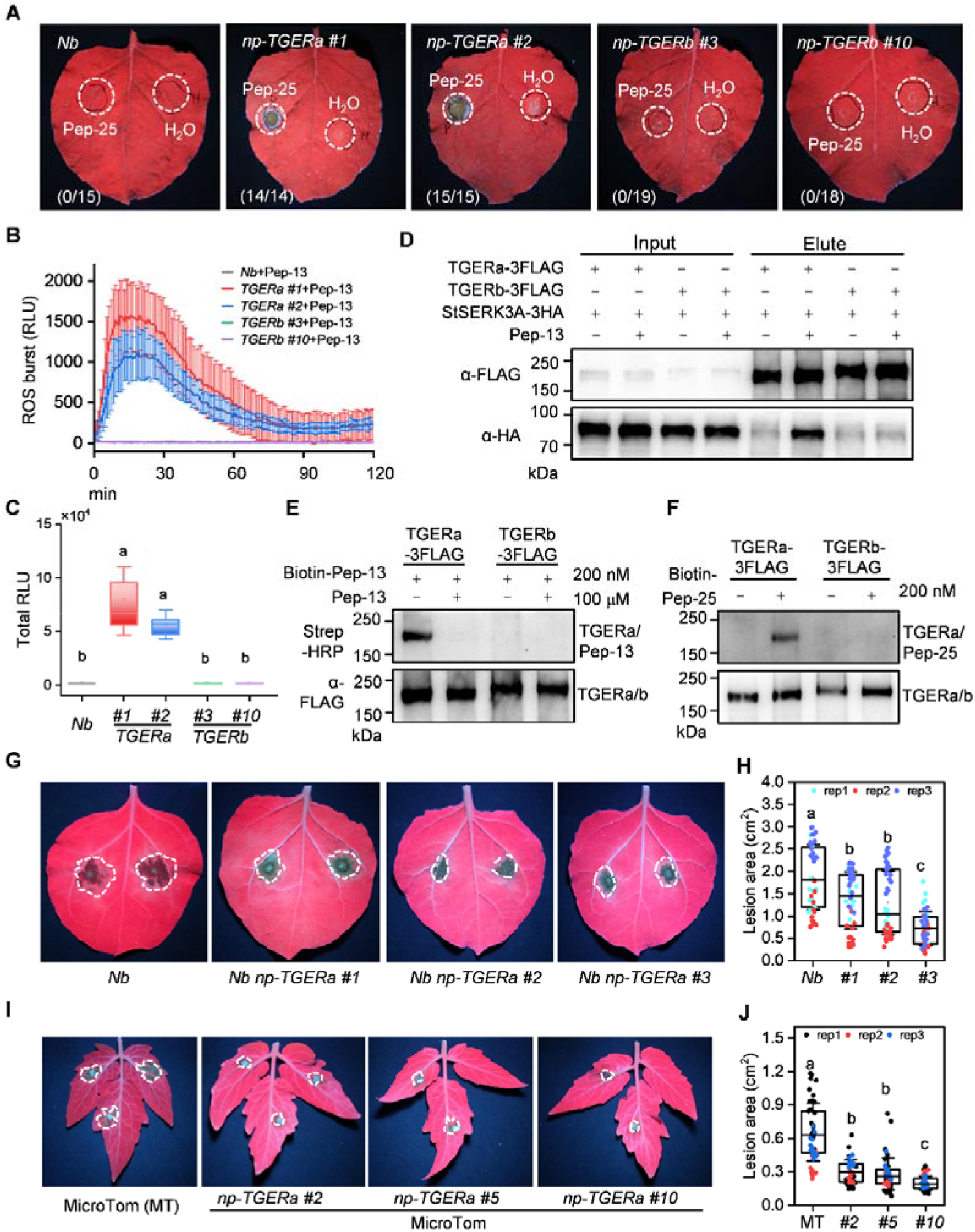
TGERa/PERU recognizes Pep-13/25 and enhances plant resistance to *Phytophthora*. (A) Pep-25-induced cell death in transgenic *N. benthamiana* (*Nb*) lines expressing *TGERa* but not *TGERb*. Leaves of transgenic lines *np-TGERa #1, #2* and *np-TGERb #3, #10* were infiltrated with 100 μM Pep-25. *Nb* plants served as a negative control. Photographs were taken under UV light 5 days after infiltration. (B-C) Pep-13-induced ROS burst in *np-TGERa* lines, but not in *np-TGERb* lines and *Nb* plants, as shown by the ROS burst curves (B) and total ROS production (C) induced by 100 nM Pep-13. Data was analyzed by two-way ANOVA followed by Tukey’s multiple comparison test. Different letters indicate significant differences (*p* < 0.05, n = 8). (D) TGERa, but not TGERb, associates with the co-receptor StSERK3A upon Pep-13 treatment. *TGERa/b-3Flag* was co-expressed with *StSERK3A-3HA* in *Nb* leaves via Agrobacterium-mediated transient expression. After 24 h, leaves were infiltrated with 100 μM Pep-13 or H□O. Total proteins were extracted 15 min later, and TGERa/b-3Flag was immunoprecipitated with M2 beads. IP products were analyzed by western blotting using anti-Flag and anti-HA antibodies. (E-F) TGERa, but not TGERb, binds biotin-labeled Pep-13 (E) and Pep-25 (F). *TGERa/b-3Flag* were transiently expressed in *Nb* leaves for 24 h. Leaves were co-infiltrated with 200 nM biotin-Pep-13/25 and 3 mM EGS, then incubated for 30 min at room temperature. Excess unlabeled Pep-13 (100 μM) was included as a competitor. TGERa/b-3Flag proteins were enriched using M2 beads, and IP eluates were detected by western blot using an anti-Flag antibody and Strep-HRP. (G-H) Transgenic *Nb* lines expressing *TGERa* exhibit enhanced resistance to *P. infestans*. (G) Detached leaves of transgenic *Nb* lines (*np-TGERa #1, #2, #3*) and *Nb* plants were inoculated with *P. infestans* strain 1306 spore suspension and photographed under UV light 5 days later. (H) Lesion areas were quantified using ImageJ. Data were analyzed using one-way ANOVA followed by Tukey’s test. Different letters (a, b, c) indicate significant differences (*p* < 0.05, n = 46 from three independent experiments). (I-J) Transgenic MicroTom tomato lines expressing *TGERa* also exhibited improved resistance to *P. infestans*. (I) Detached leaves were inoculated with *P. infestans* 1306 and photographed under UV light 2 days later. (J) Lesion areas were quantified using ImageJ. Statistical significance was determined by one-way ANOVA with Tukey’s test (*p* < 0.05, n = 45 from three independent experiments).

To explore the genetic basis of Pep-13 insensitivity in A6-10, we sequenced, assembled, and annotated its genome (Supplemental Figure 10; see details in Supplemental Note 2). The predicted TGERa^A6-10^ protein shares 98.9% amino acid identity with TGERa^E4-63^. We cloned the CDS of *TGERa*^*A6-10*^ and transiently expressed in *Nb* leaves. Similar to TGERa^E4-63^, TGERa^A6-10^ induced MAPK activation and cell death upon Pep-13 treatment (Supplemental Figure 11A-B), indicating that the TGERa^A6-10^ protein is functionally intact. Using TGERa-specific primers for ePCR in TBtools-II (Chen et al., 2023), we identified the *TGERa* locus in A6-10 genome and detected a ∼3-kb insertion within its first intron (Supplemental Figure 11C). This insertion was also present in the founder line PG6359 (Supplemental Table 4), suggesting that it may attenuate *TGERa* expression in A6-10. In line with this hypothesis, transcriptional analysis showed that *TGERa* was strongly induced in A6-26 after Pep-13 treatment, but only weakly induced in A6-10 (Supplemental Figure 11D). Furthermore, four Pep-13-responsive genes (Nietzschmann et al., 2019), namely *StTHT, StPALs, StFAD*, and *St4-CL*, were markedly upregulated in A6-26, but showed little or no induction in A6-10 (Supplemental Figure 11E). Given that Pep-25 exhibits stronger cell-death-inducing activity than Pep-13 (Supplemental Figure 3A), we further compared the responses of A6-10, A6-26, and Atlantic to both peptides. A6-26 developed robust cell death as early as 1 day after infiltration with either Pep-13 or Pep-25 (Supplemental Figure 11F-G). By contrast, Pep-25 induced delayed cell death in A6-10, while Pep-13 failed to trigger visible cell death. Neither peptide elicited cell death in Atlantic. Collectively, these results demonstrate that *TGERa*^*A6-10*^ is a weak allele with reduced expression, most likely caused by the ∼3-kb insertion in its first intron.

To assess the role of TGERa in resistance against *Phytophthora* pathogens, detached *Nb* leaves expressing empty vector (EV), *35S:TGERa*, or *35S:TGERb* were inoculated with *P. capsici* BYA5, quantification of diseased leaf areas revealed that TGERa, but not TGERb, significantly enhanced resistance to *P. capsici* BYA5 compared with EV (Supplemental Figure 9D-E). Furthermore, we challenged *np-TGERa Nb* lines with *P. infestans* 1306. Quantification of diseased leaf areas showed significantly smaller lesions in *np-TGERa* lines than in WT (Figure 2G-H). Similarly, transgenic *np-TGERa* MicroTom plants exhibited significantly reduced disease symptoms relative to MicroTom (Figures 2I-J). Collectively, these results demonstrated that introduction of *TGERa* into heterologous plant species significantly enhances their resistance against *Phytophthora* pathogens.

In summary, we identified TGERa - an allelic variant of PERU - as the receptor for Pep-13 in diploid potato inbred lines, and confirmed it mediates Pep-13 perception by forming a complex with SERK family proteins. TGERb, a close paralog of TGERa, fails to recognize Pep-13 due to its defect in ligand binding. Meanwhile, *TGERa*^*A6-10*^ is a weak allele with reduced expression and this low expression results in no visible response of A6-10 to Pep-13. We further showed that the TGERa isoform derived from intron 2 retention is non-functional; therefore, the extra 24-amino-acid peptide in the annotated PERU^LPH^ is likely annotation error rather than a determinant of broad ligand recognition. Most importantly, heterologous expression of *TGERa* in *Nicotiana benthamiana* and tomato significantly enhanced their resistance against *Phytophthora* pathogens, rendering *TGERa* a valuable candidate gene for improving *Phytophthora* resistance in other crop species.

## DATA AVAILABILITY

The authors hereby declare that all relevant data supporting the findings of this study are accessible within the research article and its supplemental information. The A6-10 genome data has been deposited at http://solomics.agis.org.cn/potato/species, under Accession A6-10. Materials generated during the current study are available from the corresponding author upon reasonable request.

## ACKNOWLEDGMENTS

We are grateful to Sanwen Huang and Chunzhi Zhang for the potato inbred lines. We kindly thank Yan Wang for pTRV2-NbSERKs, and Fanjiang Kong for pTF101-35S-3FLAG. We also acknowledge Xili Liu for sharing the *Phytophthora capsici* strain BYA5. The authors declare no conflicts of interest.

## AUTHOR CONTRIBUTIONS

T.S., X.F.,and D.L. conceived and designed this study; X.F., D.L., Y.Z., and Y.H. performed the experiments; L.C. analyzed BSA-seq data and performed assembly and annotation of the A6-10 genome. T.S., X.F., and L.C. wrote the manuscript from the input from all other authors. All authors reviewed the manuscript.

## SUPPLEMENTAL INFORMATION

Supplemental information is available Online.

