## Supplemental information for "Re-Discovery of the *Phytophthora* PAMP Pep-13 Receptor Using Potato Inbred Lines"

**Supplemental Note 1. Mapping the *TGER* locus by BSA-seq and validation.**

**Supplemental Note 2. A6-10 genome assembly and annotation.**

**Supplemental Methods**

**Supplemental Figures 1-11**

Supplemental Figure 1. Mapping the *TGER* locus using bulked segregant analysis sequencing (BSA-seq).
Supplemental Figure 2. Screening of selected candidate genes for Pep-13 response.
Supplemental Figure 3. Functional validation of TGERa by transgenic complementation.
Supplemental Figure 4. IP-MS identification of six NbSERK proteins as Pep-13-induced TGERa interactors and phylogenetic analysis of SERK family proteins.
Supplemental Figure 5. TGERa relies on SERKs for Pep-13 perception.
Supplemental Figure 6. Translation from uORF1 reduces TGERa protein accumulation.
Supplemental Figure 7. Intron 2 retention occurs in a subset of *TGERa/PERU* transcripts.
Supplemental Figure 8. TGERa-In2R derived from intron 2-retained transcripts fails to recognize Pep-13/25.
Supplemental Figure 9. TGERa, but not TGERb, mediates MAPK activation and cell death in response to Pep-13, and confers enhanced resistance.
Supplemental Figure 10. Genome assembly and annotation of potato inbred line A6-10.
Supplemental Figure 11. *TGERa* in A6-10 is a weak allele with reduced expression.

**Supplemental Tables 1-5**

Supplemental Table 1. BSA-seq-identified 1981 candidate variants on E4-63 chromosome 3.
Supplemental Table 2. SNPs identified in selected 24 candidate genes.
Supplemental Table 3. Selected candidate genes and cloning primers used.
Supplemental Table 4. ePCR results of *TGERa* primers in multiple potato genomes: A6-10, A6-26, E4-63, E86-69, PG6359, DM and Atlantic.
Supplemental Table 5. Primers used in this study.

### Re-Discovery of the *Phytophthora* PAMP Pep-13 Receptor Using Potato Inbred Lines

Xuefeng Fan[1,4], Dongyue Li[1,4], Lin Cheng[2,4], Yuqi Zhu[1,3], Yue Han[1] and Tongjun Sun[1]*

[1]Shenzhen Branch, Guangdong Laboratory for Lingnan Modern Agriculture, Genome Analysis Laboratory of the Ministry of Agriculture and Rural Affairs, Agricultural Genomics Institute at Shenzhen, Chinese Academy of Agricultural Sciences, Shenzhen, China

[2]National Key Laboratory of Tropical Crop Breeding, Shenzhen Branch, Guangdong Laboratory of Lingnan Modern Agriculture, Genome Analysis Laboratory of the Ministry of Agriculture and Rural Affairs, Agricultural Genomics Institute at Shenzhen, Chinese Academy of Agricultural Sciences, Shenzhen, China

[3]State Key Laboratory of Crop Stress Adaptation and Improvement, The Zhongzhou Laboratory for Integrative Biology, School of Life Sciences, Henan University, Kaifeng, China

[4]These authors contributed equally to this article.

**Supplemental Note 1. Mapping the *TGER* locus by BSA-seq and validation.**

**Bulked segregant analysis sequencing (BSA-seq)**

Twenty-four Pep-13 responsive (E4-63-like) and 24 non-responsive (A6-10-like) plants were selected from the $F_2$ generation of a cross between A6-10 and E4-63, respectively. Total genomic DNA was extracted and equal amounts of DNA from 24 plants were mixed to construct the E4-63-like pools and A6-10-like pools, respectively. The DNA libraries were constructed and sequenced on an Illumina platform, generating 150 bp paired-end reads. After sequencing, quality control and data preprocessing were conducted using fastp (Chen, 2025; Chen et al., 2018) to obtain clean data. These procedures were completed by Novogene Company. Whole-genome resequencing of the E4-63-like pools and A6-10-like pools generated approximately 16 GB data for each sample.

High-quality clean reads of each accessions were aligned to the E4-63 reference genome using BWA v0.7.17-r1188 (Li and Durbin, 2010) with default parameters to produce SAM file. The SAM files were then converted to BAM format with "sort" option in Samtools v1.10 (Danecek et al., 2021). The DeepVariant v1.0.0 (Shafin et al., 2021) was used to calling variation from BAM files to generate a raw variation calling format (VCF) file. This VCF file was further filtering based on AD ≥10 and annotated with SnpEff v4.5 (Cingolani et al., 2012) to predict the effects of SNPs. MutMap v2.3.2 (Sugihara et al., 2022) was run for two datasets (A6-10-like and E4-63-like) based on BAM file and VCF, using the option "-n 24" as both mutant bulks contained 24 individuals. The remaining parameters of MutMap v2.3.2 were set to default values. E4-63 was used as the reference genome, for both datasets.

The mean coverage depth for the two $F_2$ progeny pools (E4-63-like bulk and A6-10-like bulk) was approximately 21×. The genome information of E4-63 has been previously reported (http://solomics.agis.org.cn/potato/,under accession E4-63), while the genome of A6-10 was *de novo* assembled in this study (Supplemental Note 2). When the genome sequences of A6-10 were aligned to the E4-63 reference genome, a total of 11,924,729 variants, including single-nucleotide polymorphisms (SNPs) and small insertions/deletions (indels), were identified. Following validation against the parental genotypes of A6-10, E4-63 and A6-26 (http://solomics.agis.org.cn/ potato/, under accession A6-26), this number was reduced to 334,172 variants, which were further filtered based on the genotypes of the two bulked pools. Only variants that were homozygous alternative (1/1) in the A6-10-like pool and either homozygous reference (0/0) or heterozygous (0/1) in the E4-63-like pool were retained, yielding 23,142 high-quality variants. These filtered variants were subsequently subjected to SNP-index analysis (Supplemental Figure 1D).

Δ(SNP-Index) was employed to identify candidate quantitative trait locus (QTL) regions. A significant peak ($p < 0.01$) in the SNP-Index distribution was detected, spanning the 0–3 Mb region on chromosome 3 (Supplemental Figure 1E). Additionally, we identified 1,981 candidate variants with a Δ(SNP-Index) value greater than 0.99; these variants were annotated to 197 genes (Supplemental Table 1). Among these candidate genes, receptor-like proteins/receptor-like kinases (RLPs/RLKs) were prioritized as key candidates, and 24 candidate genes were selected (Supplemental Table 2 and 3). Their distribution on Chr 3 was shown (Supplemental Figure 1F).

**Candidate genes screening and validation of *TGERa***

We successfully cloned 21 of these 24 candidate genes and screened their responsiveness to Pep-13 via *Agrobacterium*-mediated transient expression in *N. benthamiana* (*Nb*) leaves. The gene expressed by construct #9 was found to respond to Pep-13 (Supplemental Figure 2), which was designated as *TGERa*. Next, we generated transgenic *Nb* lines expressing *TGERa* under its native promoter (*np-TGERa* lines); these transgenic lines exhibited cell death in response to Pep-25 (a 25-amino-acid peptide containing the Pep-13 peptide) but showed no or weak cell death to Pep-13 treatment (Supplemental Figure 3A). However, both Pep-13 and Pep-25 were able to induce MAPK activation in *np-TGERa* lines (Supplemental Figure 3B); furthermore, Pep-13 treatment of these transgenic plants significantly activated the expression of *NbACRE31* and *NbCYP71D20* (Supplemental Figure 3C). These results indicate that *TGERa* is responsive to both Pep-13 and Pep-25, with Pep-25 exhibiting stronger activity in inducing cell death than Pep-13. Stable expression of *TGERa* in tomato *cv.* MicroTOM and potato *cv.* Atlantic conferred the ability to recognize Pep-13 and Pep-25, leading to the induction of cell death (Supplemental Figure 3D-3G). Collectively, these findings demonstrated that TGERa is responsive to Pep-13/25.

**Supplemental Note 2. A6-10 genome assembly and annotation.**

The size of A6-10 genome was estimated to be ~768.3 Mb with a heterozygosity of 0.07% (Supplemental Figure 10A). For the initial contig-level assembly, we used ~22.6 Gb (~29×) of accurate circular consensus sequencing (CCS) reads, and 131.4 Gb (~171×) of high-throughput chromosome conformation capture (Hi-C) data, resulting in an assembly of 793.2 Mb (Supplemental Figure 10B). The initial assembly had a contig N50 of 27.1 Mb and was subsequently scaffolded and ordered using Hi-C data (Supplemental Figure 10C), with 99.1% of the assembled sequences anchored to chromosomes (Supplemental Figure 10B). The sequences were then deduplicated at 80% sequence similarity, and contigs shorter than 50 kb were removed. The Benchmarking Universal Single-Copy Orthologs (BUSCO) (Simao et al., 2015) was applied to evaluate the assembly completeness by identifying a set of highly conserved orthologs in the assembly. A total of 1,576 (99.5%) and 1,571 (97.3%) BUSCO genes were identified in the genome and annotation (Supplemental Figure 10B), respectively, indicating the high completeness of the A6-10 genome assembly and annotation.

**Sequencing and *de novo* genome assembly.**

The genome sequences of A6-10 were generated using the ccs program version 6.4.0 (https://github.com/PacificBiosciences/ccs) based subreads from the Pacific Biosciences Sequel II platform, which then converted to FASTQ format using SAMtools v1.17 (Danecek et al., 2021). In total, ~22.6 Gb of HiFi data were generated, representing approximately 29-fold genome coverage. For Hi-C libraries construction, DNA was extracted from seedlings and digested with the restriction enzyme MboI following previously described protocols (Burton et al., 2013; Kaplan and Dekker, 2013). A total of 131.4 Gb of Hi-C data were generated on the Illumina HiSeq platform, providing approximately 171-fold genome coverage. To facilitate genome annotation and gene expression analyses, we collected RNA-seq data from six accessions (including tissues of roots, stems, leaves, stolon, tubers and flowers) from a previous study (Tang et al., 2022). Genome assembly was performed using hifiasm v0.16 (Cheng et al., 2021) with default parameters, combining HiFi and Hi-C reads (https://github.com/chhylp123/hifiasm). The primary contigs produced by hifiasm were further scaffolded and ordered into chromosome-scale pseudomolecules using RagTag (Alonge et al., 2022), guided by the chromosome-level reference genome DMv6 (Pham et al., 2020).

**Annotation of repetitive elements.**

Transposable elements were identified using the Extensive de novo TE Annotator (EDTA) v2.1.0 (Ou et al., 2022), which include long terminal repeat retrotransposons (LTR), DNA transposons with terminal inverted repeat (TIR) sequences, and Helitron-like DNA transposons.

**Prediction of protein-coding genes.**

A comprehensive strategy consisting of transcript evidence, ab initio prediction, and homology alignment was applied for gene prediction. First, we aligned RNA-seq reads to assembled haplotypes used HISAT2 v2.2.1 (Kim et al., 2015) with the "--dta" parameter and then assembled by StringTie v2.2.1 (Pertea et al., 2015) with the "--rf" parameter. We then used BRAKER2 v2.1.5 (Bruna et al., 2021) program to train the ab initio prediction model from AUGUSTUS v3.4.0 (Stanke et al., 2006) (https://github.com/Gaius-Augustus/Augustus) and a hidden markov model (HMM) from GeneMark-ET v3.67 (Lukashin and Borodovsky, 1998) with the parameter "--nocleanup --softmasking" with high quality RNA-Seq hints. To improve the gene structure prediction, we employed a homology search method using a curated plant protein dataset downloaded from the UniProt Swiss-Prot database (https://www.uniprot.org/downloads). We merged this with previously published peptide from tomato (Sato et al., 2012) and potato (Xu et al., 2011) and eliminated potential redundancy using the CD-HIT-est (v4.6.8) (Li and Godzik, 2006) program with default parameters. MAKER2 v3.01.03 (Holt and Yandell, 2011) program was used to combine the homology search, expression evidence and ab initio prediction through two rounds. Finally, we used the Mikado v2.3.4 (Venturini et al., 2018) program to identify the most useful set of transcripts from multiple transcript assemblies, which we provided to the PASA pipeline v2.5.1 (Haas et al., 2003) for comparing and updating gene structure.

To perform functional gene annotation, we utilized the InterProScan v5.34-73.0 (Jones et al., 2014) program, which identifies potential protein domains and Gene Ontology (GO) terms based on sequence signatures. We applied the following parameters to the program: "-cli -iprlookup -tsv -gotermd -appl Pfam". In particular, we extracted protein domain information from Pfam by enabling "-appl Pfam" parameter. For each of the respective genes, we assigned the functional description of the best hit.

#### Supplemental Methods

##### Plant materials and growth conditions

The diploid potato inbred lines A6-10, E4-63, and A6-26 were previously reported (Zhang et al., 2021). Potato plants and transgenic lines were propagated on Murashige and Skoog (MS) medium in growth chambers under long-day conditions (16 h light at 23°C and 8 h dark at 22°C).

For genetic analysis, the Pep-13–insensitive line A6-10 (female parent) was artificially cross-pollinated with E4-63 (male parent) to generate $F_1$ hybrid progeny. The $F_1$ plants were subsequently self-pollinated to produce the $F_2$ population. For soil-grown seedlings, *in vitro*-cultured potato plants were grown on MS medium for 1 week and then transferred to sterile soil for further cultivation for 14 days. *N. benthamiana* and *S. lycopersicum* cv. MicroTom were directly seed-sown in soil composed of nutrient soil, vermiculite, and perlite (3:2:1, v/v/v). Plants were grown in a growth chamber under long-day conditions (16 h light at 23°C and 8 h dark at 22°C).

##### Peptide treatments

Peptides Pep-13 (VWNQPVRGFKVYE), Pep-13W2A (VANQPVRGFKVYE), and Pep-25 (DVTAGAEVWNQPVRGFKVYEQTKMS) (Brunner et al., 2002), Biotin-Pep-13, Biotin-Pep-25 and flg22 (QRLSTGSRINSAKDDAAGLQIA) were synthesized by Sangon Biotech (Shanghai, China). Peptides were dissolved in sterile double-distilled water ($ddH_2O$) to generate 2 mM stock solutions and stored at -20°C. Working solutions were prepared by diluting stock solutions with sterile water to indicated concentrations.

##### Vector construction

Primers used in this study are listed in Supplemental Table 5. The full-length coding sequences of *TGERa*, *TGERb*, *StSERK2* and *StSERK3A*/*B* were cloned from the cDNA library derived from of E4-63 leaf. PCR fragments were inserted into pTF101-35S-3Flag vector (Bu et al., 2021) digested with *XbaI* and *MluI* via homologous recombination. *StSERK2* and *StSERK3A*/*B* were inserted into pCAMBIA1300-35S-3HA vector digested with *HindIII* and *KpnI* via homologous recombination. All constructs were verified by Sanger sequencing.

##### *Agrobacterium*-mediated transient expression

Recombinant *Agrobacterium tumefaciens* strain GV3101 harboring the indicated constructs was cultured overnight in LB medium supplemented with kanamycin (50 μg/mL), rifampicin (25 μg/mL), and gentamycin (50 μg/mL) at 28 °C with shaking at 200 rpm. Bacterial cells were harvested by centrifuged at 5000 rpm for 5 min and washed twice with the infiltration buffer (10 mM $MgCl_2$, 10 mM MES (pH 5.6), and 200 μM acetosyringone). Cells were resuspended in infiltration buffer to a final $OD_{600}$ of 0.6 and incubated at room temperature for 3 h. Fully expanded leaves of 4-5-week-old *N. benthamiana* plants were infiltrated on the abaxial side using 1-ml syringe. Plants were maintained under low light conditions for 24 h and then returned to normal growth conditions. Protein expression was verified by immunoblot analysis 24-48 h after infiltration. For elicitor assays, 100 μM Pep-13 was infiltrated at 24 h post infiltration. Hypersensitive cell death was evaluated 48 h later under 365 nm UV light.

**RNA isolation and RT-qPCR**

RNA isolation and RT-qPCR was performed as described (Chen et al., 2024). *NbEF1α* was used as the reference gene for *N. benthamiana* samples, and *StTUB* was used as the reference gene for potato. Relative expression levels were calculated using the $2^{-\Delta\Delta CT}$ method. Primer sequences are listed in Supplemental Table 5.

**MAPK activation assays**

*N. benthamiana* leaves were infiltrated with recombinant *Agrobacterium tumefaciens* GV3101 and incubated for 24 hours as described previously. Subsequently, the leaves were infiltrated with 50 uM peptides (Pep-13, Pep-25, W2A) and incubated for 15 min. Proteins were separated by 10% SDS-PAGE, and the target proteins were detected using anti-p44/42-ERK antibody (Cell Signaling #9101).

**Virus-induced gene silencing (VIGS)**

Cotyledons of 2-week-old *N. benthamiana* plants or transgenic *np-TGERa* lines were infiltrated with a 1:1 mixture of *Agrobacterium* cultures containing pTRV1 and pTRV2 derivatives, *pTRV2-NbPDS*, *TRV2-GFP*, and *pTRV2-NbSERKs* (Wang et al., 2018) adjusted to $OD_{600}$ = 0.5. Two weeks post-infiltration, plants showing photobleaching in *pTRV2-NbPDS*. Gene silencing efficiency was confirmed by RT-qPCR. Leaves were then treated with 100 μM Pep-25 or $H_2O$ and cell death responses were documented under 365 nm UV light 48 h later.

**Electrolyte leakage assay**

Electrolyte leakage assay was performed as reported (Fan et al., 2026). Briefly, leaf discs (8 mm) were punched from peptide- or water-infiltrated leaves. Six discs per replicate immersed in 2 mL deionized water. After 3 h incubation at 25 °C, initial conductivity (R1) was measured. Samples were boiled for 30 min, cooled to 25 °C, and final conductivity (R2) recorded. Relative electrolyte leakage (REL) was calculated as: REL (%) = (R1/R2) × 100%.

**Stable transformation of *N. benthamiana*, *Solanum tuberosum* (cv. Atlantic) and *S. lycopersicum* (cv. MicroTom)**

Stable transformation of *N. benthamiana*, MicroTom and Atlantic plants was carried out by the BioRun company via *Agrobacterium*-mediated transformation, using the construct *np-TGERa-3Flag* for *N. benthamiana* and MicroTom, and the construct *35S-TGERa* for Atlantic. Transgenic plants were screened by PCR using gene-specific primer to confirm the presence of the *TGERa* transgene. TGERa protein levels in positive lines were further verified by Western blot analysis.

***In vivo* ligand binding assays**

*Agrobacterium tumefaciens* GV3101 carrying *TGERa-3flag* or *TGERb-3flag* constructs was transiently expressed in *N. benthamiana* leaves for 24 h. Leaves were then infiltrated with solutions containing 100 nM biotin-Pep-13/25 alone or together with 100 μM unlabeled Pep13/25 as a competition. The infiltration buffer contained 3 mM ethylene glycol bis (succinimidyl succinate) (EGS，Thermo Scientific, #21565) in 25 mM HEPES (pH 7.5). After 30 minutes, approximately 1 g of leaf tissue was harvested and frozen in liquid nitrogen. Proteins were enriched using anti-Flag M2 affinity resin (Sigma-Aldrich, A2220). Bound proteins were analyzed by immunoblotting using anti-Flag antibody (Sigma-Aldrich, F1804) and streptavidin-HRP (Abcam, Ab7403) to detect receptor-bound biotinylated peptides.

**Co-immunoprecipitation assay**

Frozen leaf tissue was ground into a fine powder in liquid nitrogen and homogenized in membrane extract buffer containing 25 mM Tris-HCl (pH 7.5), 10% Glycerol, 1 mM EDTA, 150 mM NaCl, 1.5% NP-40, 0.5% sodium deoxycholate, 1 mM NaF, 4 mM DTT, 2% (w/v) PVPP, 1 mM PMSF, and protease cocktail. Protein extracts were incubated on ice for 30 min and centrifuged at 13,000 × g for 15 min at 4 °C. The supernatants were clarified by a second centrifugation under the same conditions. For immunoprecipitation, the supernatants were incubated with 20 μL anti-FLAG M2 affinity resin for 2 h at 4 °C with gentle rotation. Input samples were retained for immunoblot analysis. The resin was washed three times with binding buffer containing 25 mM Tris-HCl (pH 7.5), 10% Glycerol, 1 mM EDTA, 150 mM NaCl, 1.5% NP-40, 0.5% sodium deoxycholate, 1 mM NaF, 0.2 mM DTT. Bound proteins were eluted by boiling in 2× SDS loading buffer at 95 °C for 5 min. Proteins were separated by SDS–PAGE and detected by immunoblotting using anti-Flag and anti-HA antibodies.

**Interacting proteins identified by Immunoprecipitation-Mass Spectrometry （IP-MS）**

*TGERa-3Flag* or empty vector (EV) constructs were transiently expressed in *N. benthamiana* leaves. At 48 h after agroinfiltration, leaves were treated with 100 μM Pep-13 or $H_2O$ and harvested 15 min after treatment. Total proteins were extracted and subjected to immunoprecipitation using anti-FLAG M2 affinity resin as described above. Immunoprecipitated proteins were subjected to trypsin digestion. The resulting peptides were analyzed by liquid chromatography-tandem mass spectrometry (LC-MS/MS) using an EASY-nLC 1200 system coupled to an Orbitrap Exploris 480 mass spectrometer (Thermo Fisher Scientific). Peptides were separated on a C18 analytical column using a linear acetonitrile gradient.

Raw MS data were processed using Proteome Discoverer software and searched against the NbHZ1 protein database (Wang et al., 2024). Carbamidomethylation of cysteine residues was set as a fixed modification, whereas methionine oxidation was considered a variable modification. Up to two missed trypsin cleavage sites were allowed. Peptide spectrum matches were filtered using a false discovery rate (FDR) threshold of <1%. Functional annotation of identified proteins was performed using Gene Ontology (GO), InterPro, COG, and KEGG databases. Only proteins identified in at least two biological replicates were considered for further analysis.

**Luminol-based ROS burst assay**

Leaf discs (4 mm diameter) were excised from 5-week-old *N. benthamiana* plants were equilibrated overnight in 200 μL dd$H_2O$ in 96-well white plates. The next day, water was replaced with 100 μL reaction buffer containing 10 μg/mL horseradish peroxidase (HRP; Sigma #P6782), 50 μM luminol (Sigma #A8511), and 50 nM Pep-13. Real-time luminescence was recorded at 1 min intervals for 120 min using a Cytation5 microplate reader (BioTek, Winooski, VT, USA) with the following parameters: gain 130, 5 mm read height, no optical filter, and maintaining a constant temperature of 22 °C.

**Pathogen infection assay**

Infection assays were performed as described previously (Chen et al., 2024). *P. infestans* strain 1306 was cultured on rye agar medium at 18 °C in darkness for 14 days. Zoospores were released by incubating sporangia in cold water at 4 °C for 2 h and filtered to remove mycelium. Detached leaves were inoculated with 5 μL zoospore suspension (1 $\times 10^5$ spores/mL) on the abaxial surface.

For *P. capsica* strain BYA5 (Gao et al., 2023), clones were cultured on V8 juice agar medium at 25 °C in darkness for 3 days. Agar plugs (5 mm diameter) from colony margins were placed onto leaf surface and covered with 10 μL sterile water. Infected leaves were incubated under 100% relative humidity at optimal temperatures (18 °C for *P. infestans*, 25 °C for *P. capsica*) for 4 or 3 days, respectively. Lesion areas were visualized under UV light and quantified using ImageJ (v1.51j8, National Institutes of Health, Maryland, USA) software.

**Statistical Analysis**

All experiments were performed with at least three independent biological replicates unless otherwise stated. Data are presented as mean ± standard deviation (SD). Statistical analyses were performed using OriginPro2025 (OriginLab). Statistical significance between two groups was determined using a two-tailed Student's t-test, while comparisons among multiple groups were analyzed using one-way analysis of variance (ANOVA) followed by Tukey's multiple comparison test. Differences were considered statistically significant at $p < 0.05$.

#### Supplemental Figures 1-11

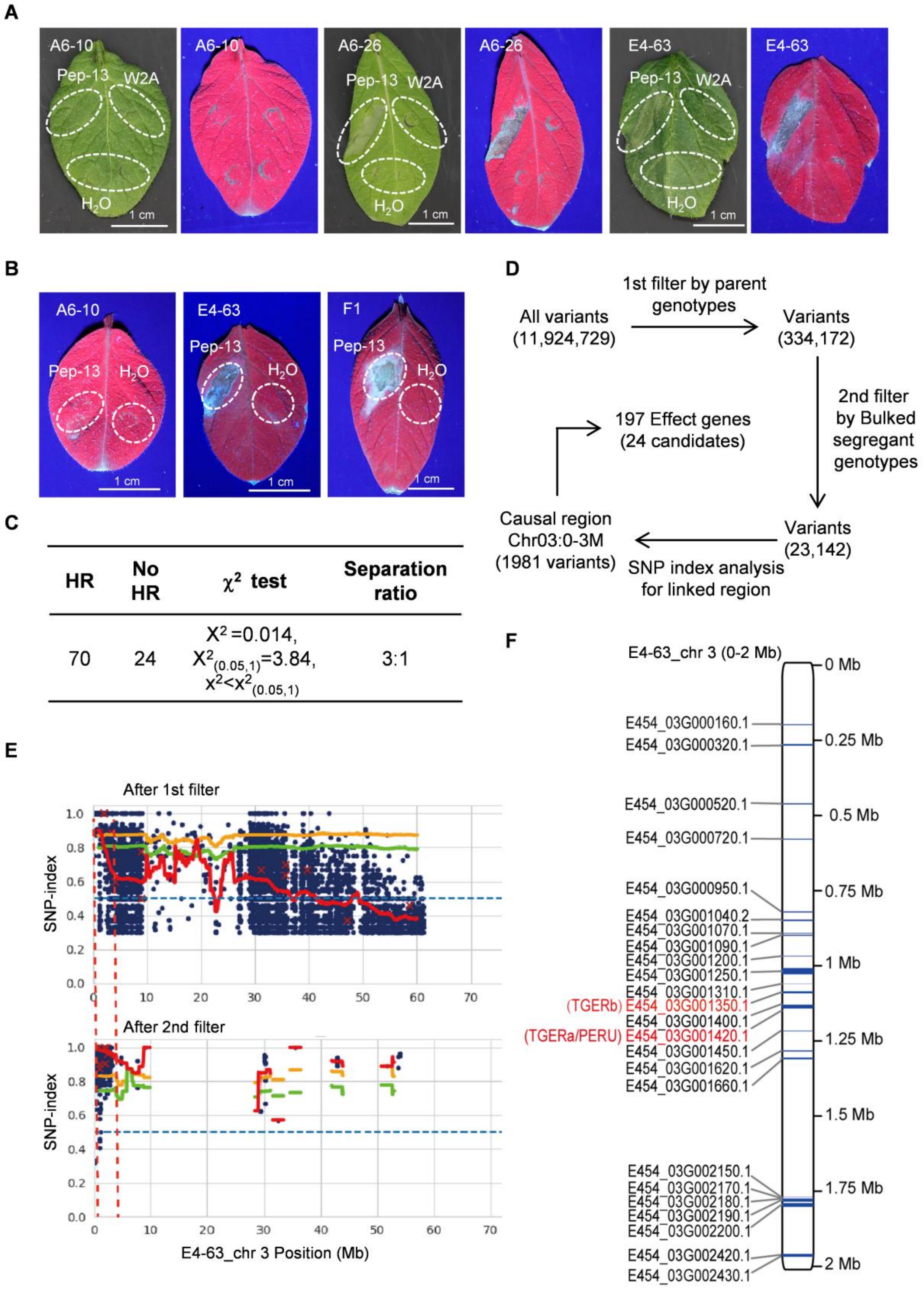


**Supplemental Figure 1. Mapping the *TGER* locus using bulked segregant analysis sequencing (BSA-seq).**

**(A)** Differential responses of potato cultivars to Pep-13. Leaves were infiltrated with 100 μM Pep-13, Pep-13W2A (an inactive Pep-13 variant) and $H_2O$. Images were taken under white light and UV light at 24 h post-infiltration. Scale bar = 1 cm..

**(B)** Responses of parental lines A6-10, E4-63 and their $F_1$ progeny to Pep-13. Leaves were infiltrated with 100 μM Pep-13 or $H_2O$. Images were captured under UV light at 48 h post-infiltration. Scale bar = 1 cm.

**(C)** Segregation of Pep-13 induced cell death in the $F_2$ population derived from self-pollinated $F_1$ plants. Leaves were infiltrated with 100 μM Pep-13, and cell death response was scored at 48 h post-infiltration.

**(D)** Workflow of the variant filtering process and the causal region identified by Δ(SNP-index) analysis.

**(E)** Manhattan plot showing the SNP-index distribution of variants on chromosome 3 after the first round of filtering (upper panel) and the second round of filtering (lower panel). Blue dots represent variants (SNPs and small Indels). The orange and green lines represent 99% and 95% confidence intervals of slide windows (200 kb window), respectively. The red line represents mean SNP-index. The horizontal blue dashed line indicates the threshold (0.5). The causal region (0-3 Mb) is marked with vertical red dashed lines. Numbers on the horizontal axis represent genome positions.

**(F)** Selected 24 candidate genes locates within the 0-2 Mb interval on chromosome 3 of the E4-63 genome.*TGERb* is identical to *E454_03G001350*. In contrast, the *TGERa*/*PERU* locus was not annotated in the E4-63 genome and overlaps with the incorrectly annotated locus *E454_03G001420*.

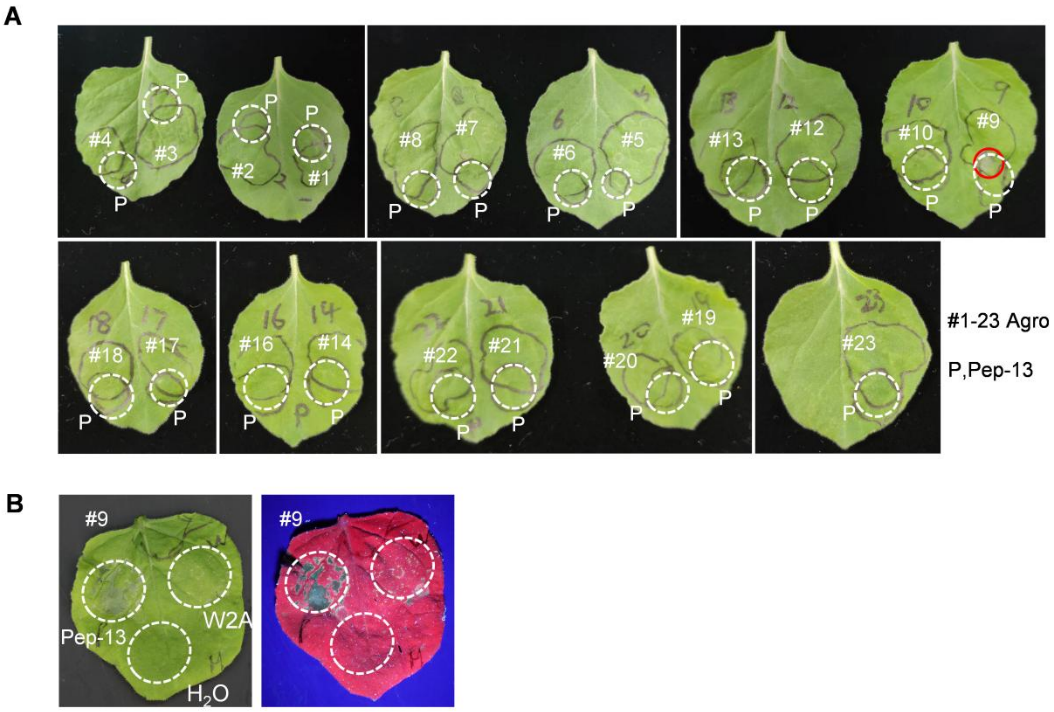


**Supplemental Figure 2. Screening of selected candidate genes for Pep-13 response.**

**(A)** Functional screening of 21 candidate genes by *Agrobacterium*-mediated transient expression in *N. benthamiana*. The corresponding cDNAs were cloned and transiently expressed in *N. benthamiana* leaves. Twenty-four hours later, leaves were infiltrated with 100 μM Pep-13 at sites overlapping the *Agrobacterium* infiltration zones (white dashed cycle). Cell death was observed specifically in the region expressing candidate gene #9 (red circle) at 24 h after Pep-13 treatment. Detailed information on these 21 candidate genes is provided in Supplemental Table 3.

**(B)** Pep-13-dependent responses mediated by candidate gene #9 (*TGERa*). *Nb* leaves transiently expressing *TGERa* were infiltrated with 100 µM Pep-13, Pep-13W2A, and $H_2O$. Cell death was observed only upon Pep-13 treatment.

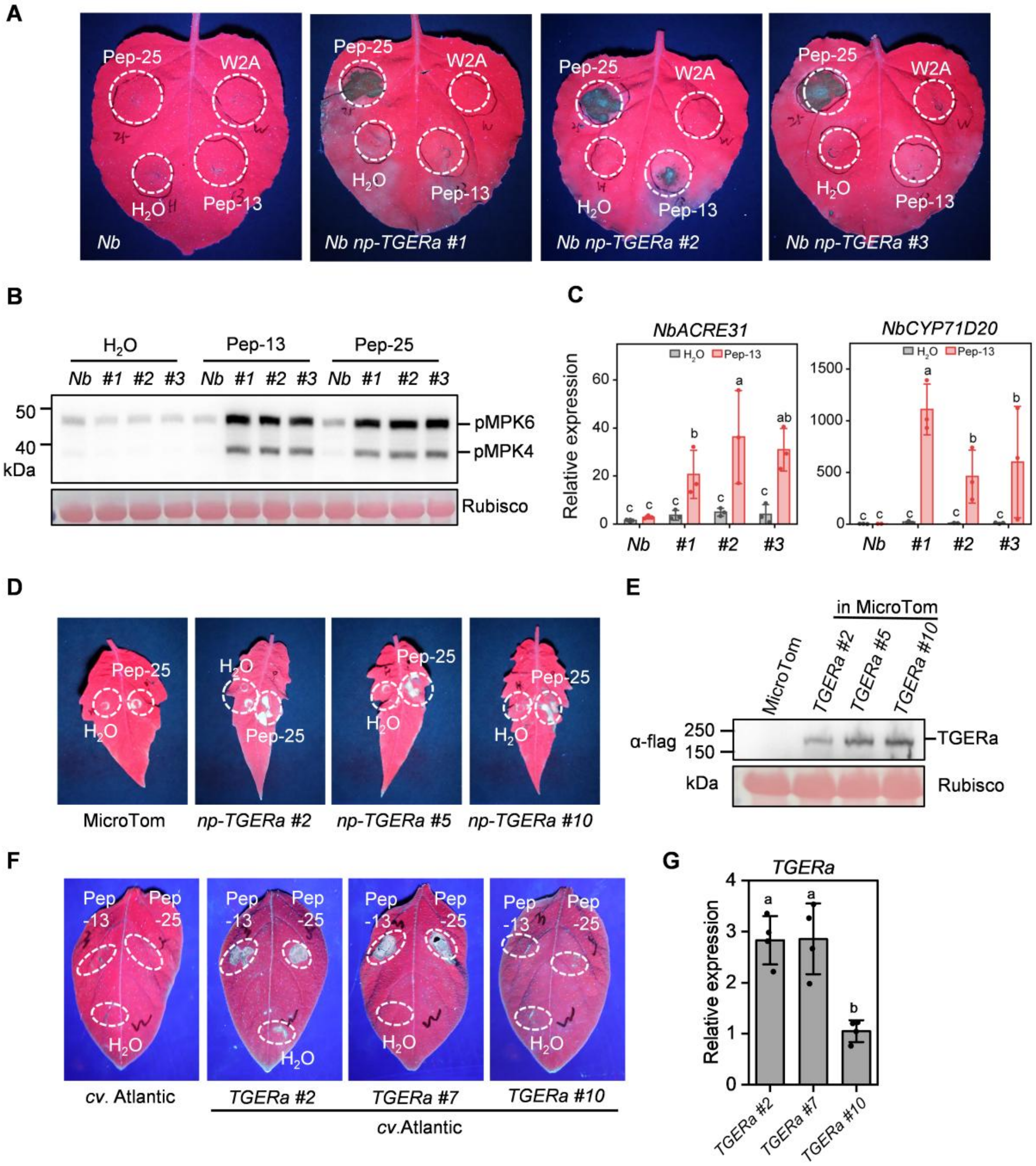


**Supplemental Figure 3. Functional validation of TGERa by transgenic complementation.**

**(A)** Cell death responses triggered by peptide treatments in transgenic *Nb* lines expressing *TGERa* under its native promoter *(np-TGERa #1*, *#2*, *#3*). Leaves were infiltrated with 100 µM Pep-25, Pep-13, Pep-13W2A or $H_2O$. *Nb* plants served as a negative control. Cell death in *np-TGERa* lines was induced 5 days after 100 µM Pep-25 treatment.

**(B)** Both Pep-13 and Pep-25 induced MAPK activation in *np-TGERa* lines. Leaves of *np-TGERa #1*, *#2*, *#3* were infiltrated with 50 µM Pep-13 or Pep-25, and MAPK samples were collected 15 min later and analyzed by western blot using anti-p44/42-ERK antibodies.

**(C)** Induction of PTI marker genes in leaves of *np-TGERa* lines. Transcript levels of *NbACRE31* and *NbCYP71D20* were measured by RT-qPCR at 4 h after infiltration with 50 µM Pep-13. *NbEF1α* used as reference gene. Statistical significance was determined by two-way ANOVA followed by Tukey's multiple comparison test. Different letters (a,b,c) indicate significant differences ($p < 0.05$, n = 3 from three independent biological replicates).

**(D)** Pep-25 induced cell death in transgenic tomato MicroTOM lines (*np-TGERa #2*, *#5*, *#10*). Leaves were infiltrated with 100 μM pep-25, and necrotic symptoms were recorded 3 days post-infiltration.

**(E)** Western blot detection of TGERa-3Flag protein levels in transgenic MicroTom lines described in (D).

**(F)** Hypersensitive cell death induced by 100 μM Pep-13 and 100 μM Pep-25 in transgenic potato Atlantic lines expressing *TGERa*. $H_2O$ treatment served as a negative control.

**(G)** Relative expression levels of *TGERa* in transgenic Atlantic lines shown in (F) determined by RT-qPCR. *StTUB* used as reference gene. Data were analyzed using two-way ANOVA followed by Tukey's multiple comparison test. Different letters (a,b) indicate significant differences ($p < 0.05$, n = 3 from three independent biological replicates).

**A**

| IP-MS experiment | Replicate 1 (230511) | | | Replicate 2 (230627) | | | | |
|---|---|---|---|---|---|---|---|---|
| Unique peptide identified in proteins (NbHZ1 ID) | TGERa-3Flag | TGERa-3Flag +Pep-13 | Nb_EV | TGERa-3Flag | TGERa-3Flag +Pep-13 | Nb_EV | Protein name | Homologs in E4-63 ( Potato inbred line) |
| TGERa | 38 | 37 | - | 45 | 43 | - | - | TGERa |
| Nbe13g26960.1 | - | 7 | - | - | 9 | - | SERK3b (E3VXE7) | StSERK3A (E454_10G011500.2) |
| Nbe06g11390.1 | - | 7 | - | - | - | - | SERK3a (E3VXE6) | |
| Nbe07g12510.1/ Nbe08g30080.1 | - | 6 | - | - | 5 | - | SERK2a / SERK2b | StSERK2 (E454_04G026450.1) |
| Nbe02g18010.1 | - | 10 | - | - | 11 | - | NbBAK1a | StSERK3B (E454_01G040240.1) |
| Nbe01g18330.1 | - | 12 | - | - | 13 | - | NbBAK1b | |

**B**

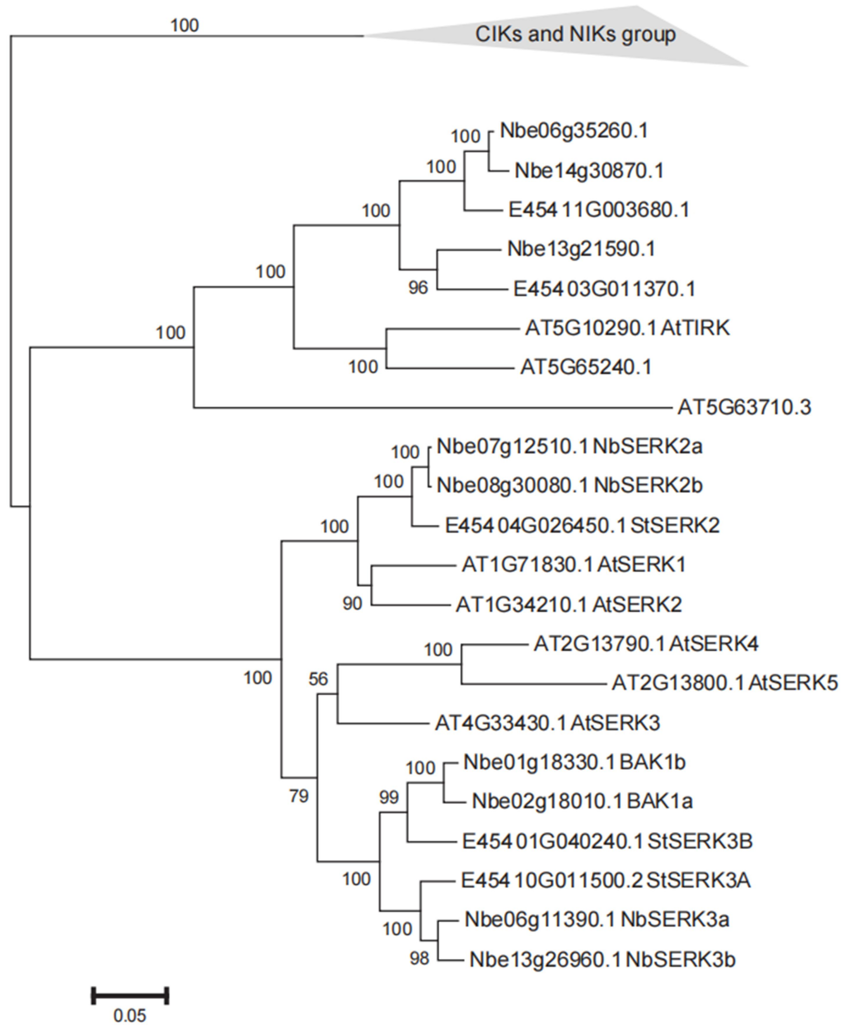


**Supplemental Figure 4. IP-MS identification of six NbSERK proteins as Pep-13-induced TGERa interactors and phylogenetic analysis of SERK family proteins.**

**(A)** Identification of TGERa-associated proteins by immunoprecipitation followed by mass spectrometry (IP-MS) in *N. benthamiana*. *TGERa-3Flag* was transiently expressed in leaves, and proteins were enriched by M2 beads in the presence or absence of Pep-13, respectively. Six NbSERK proteins, including NbSERK3a (uniprot ID E3VXE7), NbSERK3b (uniprot ID E3VXE6), NbBAK1a, NbBAK1b, NbSERK2a and NbSERK2b were identified as Pep-13 induced interacting proteins of TGERa. Three StSERK proteins, StSERK2, StSERK3A and StSERK3B were identified in E4-63 proteome by BLAST analysis.

**(B)** Phylogenetic analysis of SERK proteins from *A. thaliana*, *N. benthamiana*, and potato (E4-63). SERK protein sequences were retrieved from TAIR (https://www. arabidopsis.org), NbHZ1 genome database (http://lifenglab.hzau.edu.cn/Nicomics) and E4-63 (http:// solomics.agis.org.cn/potato/species), respectively. An unrooted maximum-likelihood phylogenetic tree was constructed with MEGA 11 software based on 1,000 bootstrap replicates. Evolutionary distances were estimated using the Poisson correction model, with branch lengths indicating the number of amino acid substitutions per site. Phylogenetic analysis revealed that immune-related SERK proteins from the three species clustered into a single clade.

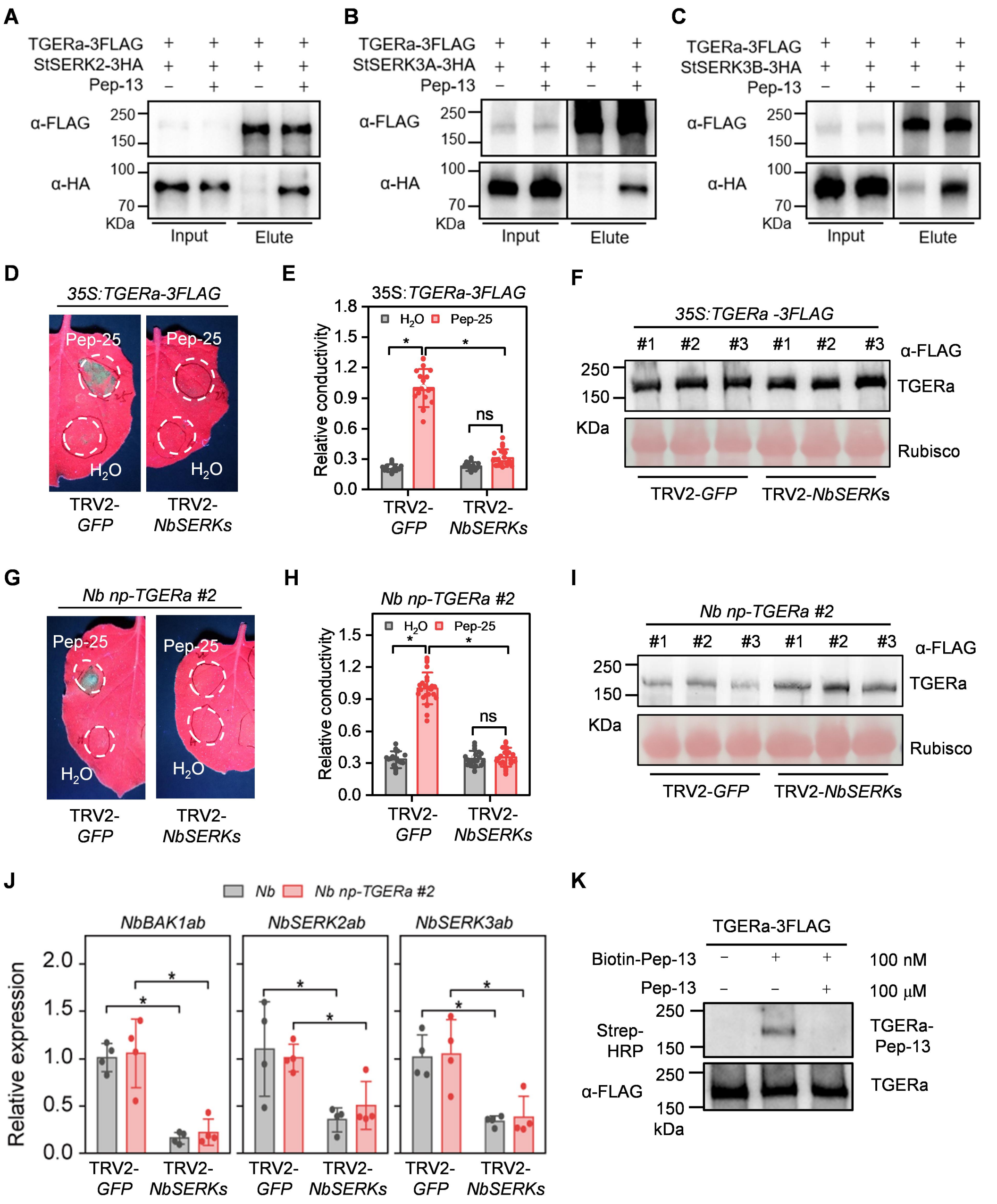


**Supplemental Figure 5. TGERa relies on SERKs for Pep-13 perception.**

**(A-C)** Co-immunoprecipitation (Co-IP) assays revealing Pep-13-induced interactions between TGERa and StSERK proteins (StSERK2, StSERK3A/B). *TGERa-3Flag* and *StSERK-3HA* were transiently co-expressed in *Nb* leaves. After 24 h, leaves were treated with 100 μM Pep-13 or $H_2O$ for 15 min before sampling. TGERa-3Flag was enriched with M2 beads, and co-precipitated StSERK-3HA was detected by western blot using anti-Flag and anti-HA antibodies.

**(D-F)** Silencing of *NbSERK* genes via VIGS impairs Pep-25-triggered hypersensitive cell death in *Nb* leaves transiently expressing *TGERa-3Flag*. (D) 100 μM Pep-25 induced cell death in plants expressing *TGERa-3Flag* in the *GFP*-silenced control, but not in *NbSERK*-silenced plants. (E) Quantification of cell death by electrolyte leakage corresponding to (D). Student's t-test, *$p < 0.05$, ns, not significant difference, n = 12 from two independent replicates. (F) Western blot analysis of TGERa-3Flag protein levels corresponding to (D).

**(G–I)** Silencing of *NbSERK* genes via VIGS impairs Pep-25-triggered hypersensitive cell death in transgenic *Nb* line *np-TGERa #2*. (G) 100 μM Pep-25 induced cell death in *GFP*-silenced *np-TGERa #2* plants, but not in *NbSERK*-silenced *np-TGERa #2* plants. (H) Quantification of cell death by electrolyte leakage corresponding to (G). Student's t-test, *$p < 0.05$, ns, not significant difference, n = 12 from two independent replicates. (I) Western blot analysis of TGERa-3Flag protein levels corresponding to (G).

**(J)** qRT-PCR analysis of silencing efficiency for *NbBAK1ab*, *NbSERK2ab*, and *NbSERK3ab* in wild-type and *np-TGERa #2* plants. Data are mean ± s.d. (n = 4); Student's t-test, *$p < 0.05$.

**(K)** Binding of TGERa to Pep-13. *TGERa-3Flag* was transiently expressed in *Nb* leaves for 24 h, Leaves were then co-infiltrated with 100 nM biotin-Pep-13 and 3 mM EGS, followed by incubation at room temperature for 30 min. Excess unlabeled Pep-13 (100 μM) was included as a competitor. TGERa-3Flag was enriched with M2 beads, and biotin-Pep-13 binding was analyzed by western blot using anti-Flag antibody and Strep-HRP.

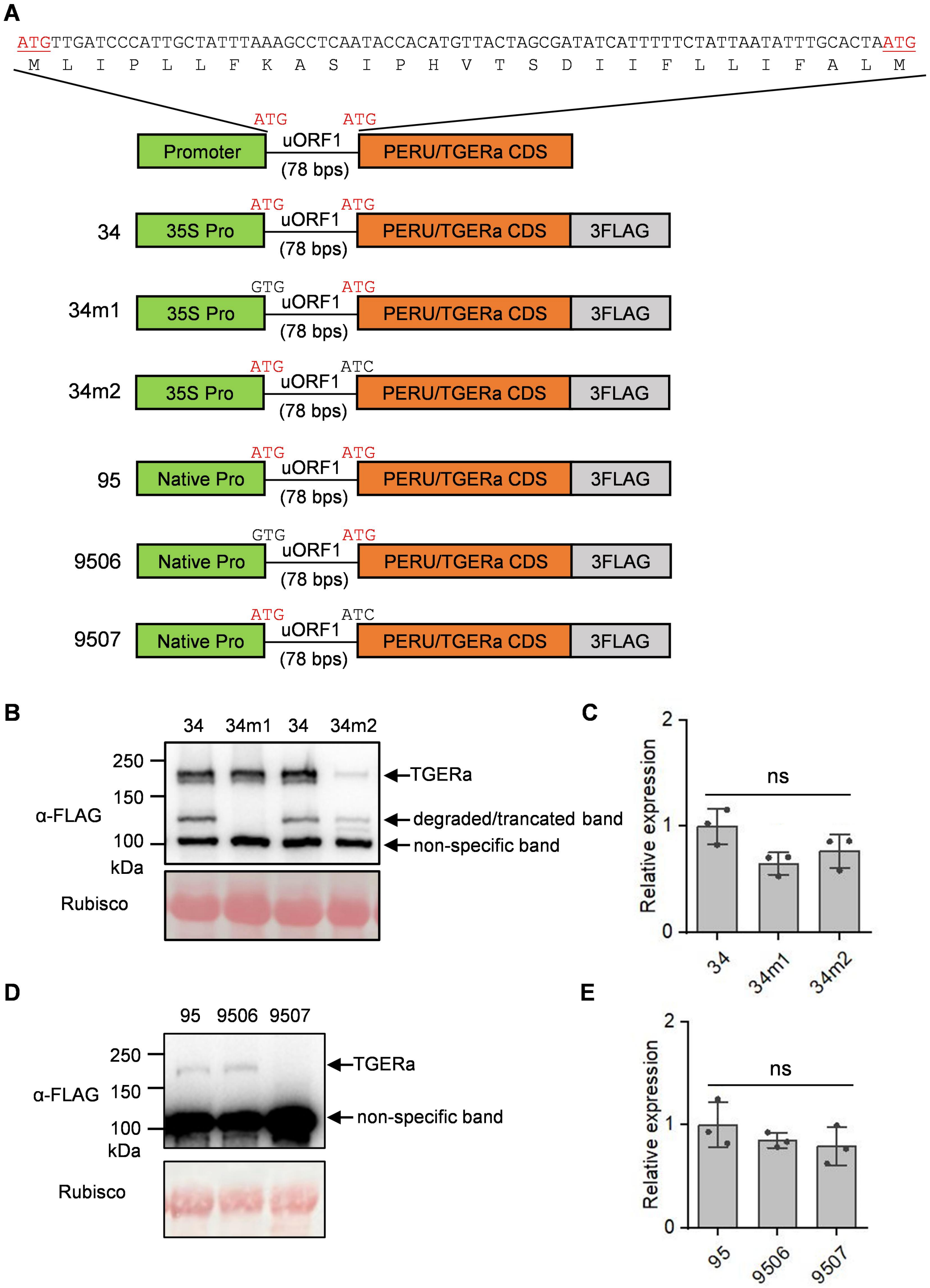


**Supplemental Figure 6. Translation from uORF1 reduces TGERa protein accumulation.**

**(A)** The uORF1 of *TGERa* mRNA is in-frame with its main ORF. Constructs 34, 34m1, and 34m2 were designed to express *TGERa* with an intact uORF1, a mutated uORF1 start codon (ATG to GTG), and a mutated main ORF start codon (ATG to ATC), respectively, under the CaMV 35S promoter. In contrast, constructs 95, 9506, and 9507 were generated to express *TGERa* and its variants under its native promoter.

**(B)** Western blot detection of TGERa-3Flag protein levels expressed from constructs 34, 34m1, and 34m2 in *Nb* leaves. Mutation in the main ORF start codon greatly reduced TGERa protein accumulation.

**(C)** Relative transcript levels of *TGERa* expressed from the 35S-driven constructs measured by qRT-PCR. n = 3, ns, no significant difference.

**(D)** Western blot detection of TGERa protein accumulation expressed from constructs 95, 9506, and 9507 in *Nb* leaves. Mutation in the main ORF start codon blocked TGERa protein accumulation under native promoter conditions.

**(E)** Relative transcript levels of *TGERa*-3Flag expressed from native promoter constructs. (B-E) Samples were collected 48 h after transient expression. Ponceau S staining of Rubisco served as a loading controls. Data represent mean ± s.d. n = 3, ns, no significant difference. Date were analyzed using Student's t-test.

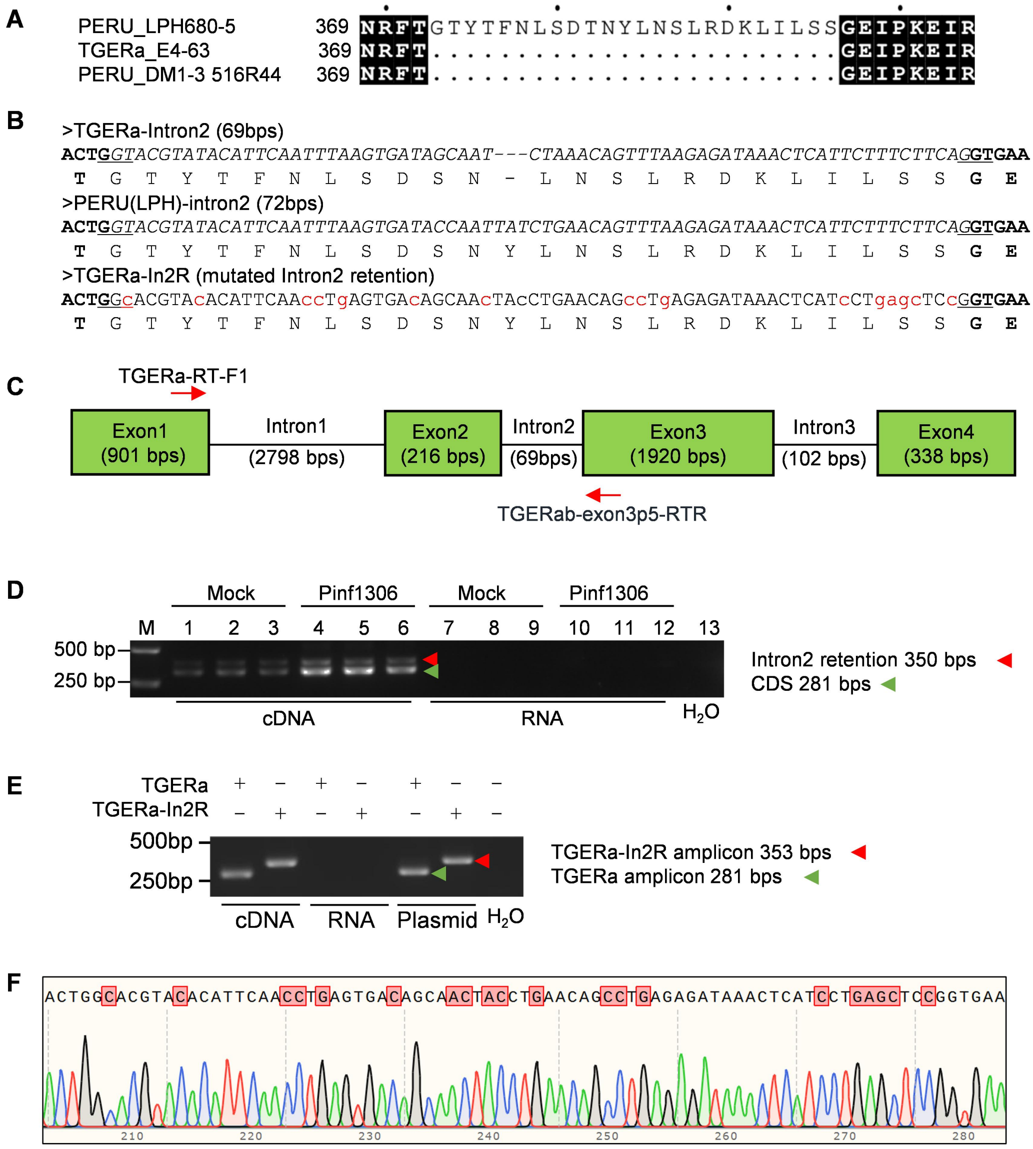


**Supplemental Figure 7. Intron 2 retention occurs in a subset of *TGERa/PERU* transcripts.**

**(A)** The annotated PERU$^{LPH680-5}$ has extra 24 amino acids compared to PERU/TGERa.

**(B)** The extra peptide is encoded by the in-frame intron 2 in the *PERU*$^{LPH680-5}$ transcript.

**(C)** Gene structure of *TGERa*, indicating the sizes of exons and introns, as well as the positions of the

RT primers used for detecting intron 2 retention. Primer sequences are listed in Supplemental Table 6. This RT primer pair spans the large intron 1 (2798bps) to avoid genomic DNA contamination.

**(D)** *TGERa* intron 2 is retained in a portion of *TGERa* mRNA extracted from A6-26 leaf tissues at 24 h after mock treatment or inoculation with *P. infestans* 1306 spore suspension. The red arrowhead indicates the cDNA fragment with intron 2 retention, whereas the green arrowhead marks the spliced cDNA fragment. Non-reverse-transcribed RNA samples served as negative controls.

**(E)** Mutation of intron 2 prevents it splicing out from *TGERa-In2R* mRNA. *TGERa* and *TGERa* with synonymous mutations in intron 2 (*TGERa-In2R,* as shown in (B)) were constructed and transiently expressed in *Nb* leaves. After 24 h, total mRNA was extracted and reverse-transcribed into cDNA; these samples were then examined by PCR and confirmed that the mutated intron 2 was retained in *TGERa-In2R* mRNA. RNA and plasmid samples served as negative and positive controls, respectively.

**(F)** Retention of the mutated intron 2 in the cDNA amplicon derived from *TGERa-In2R* mRNA in (E) was further confirmed by Sanger sequencing.

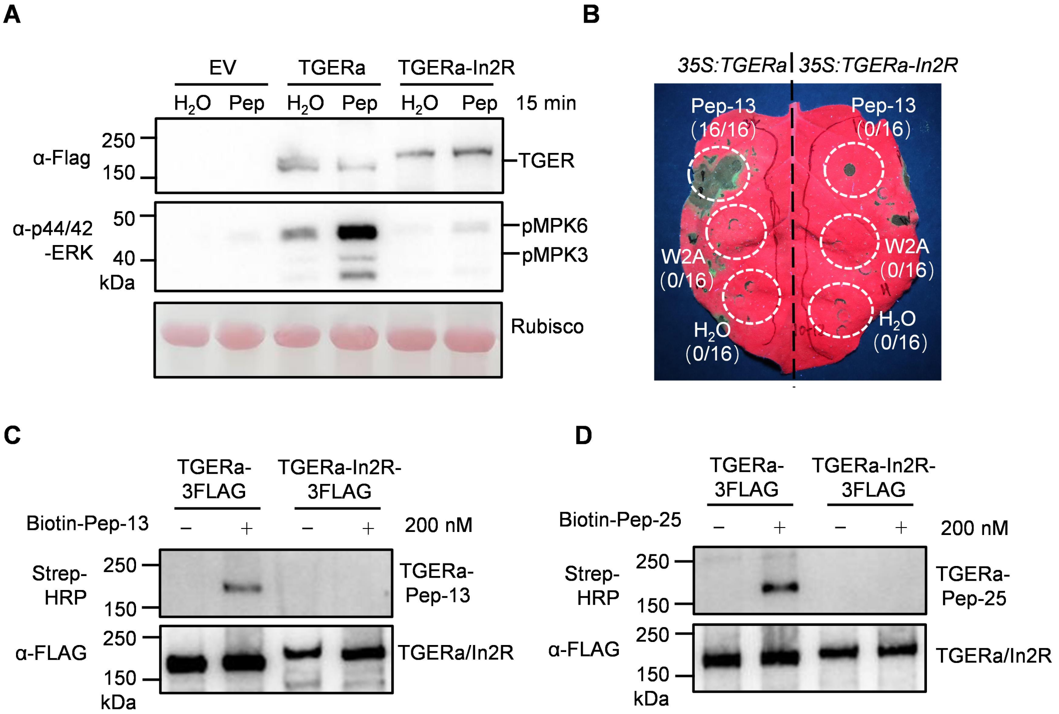


**Supplemental Figure 8. TGERa-In2R derived from intron 2-retained transcripts fails to recognize Pep-13/25.**

**(A-B)** TGERa-In2R fails to activate MAPK or induce cell death in response to Pep-13. Constructs harboring *35S:TGERa-3Flag* and *35S:TGERa-In2R-3Flag* were transiently expressed in *Nb* leaves mediated by *Agrobacterium*. After 24 h, 50 µM Pep-13 and $H_2O$ were infiltrated into leaves expressing *TGERa-3Flag* and *TGERa-In2R-3Flag*, respectively. MAPK samples were collected 15 min post-infiltration and analyzed by western blot using anti-Flag and anti-p44/42-ERK antibodies (A). For cell death analysis, 100 µM Pep-13, 100 µM Pep-13W2A and $H_2O$ were infiltrated into leaves expressing *TGERa-3Flag* and *TGERa-In2R-3Flag*, respectively. Cell death was observed only in *Nb* leaves expressing *TGERa-3Flag* after Pep-13 treatment. Numbers in parentheses show the count of necrotic spots versus total spots (B).

**(C-D)** TGERa-In2R cannot bind biotin-labeled Pep-13/25. Twenty-four hours after transient expression of *TGERa-3Flag* and *TGERa-In2R-3Flag* in *Nb* leaves, 200 nM biotin-Pep-13 (C) and biotin-Pep-25 (D) were co-infiltrated with 3 mM EGS into the leaves and incubated for 30 min at room temperature. TGERa-3Flag and TGERa-In2R-3Flag proteins were enriched using M2 beads, and the IP eluates were detected by western blot using anti-Flag antibody and Strep-HRP.

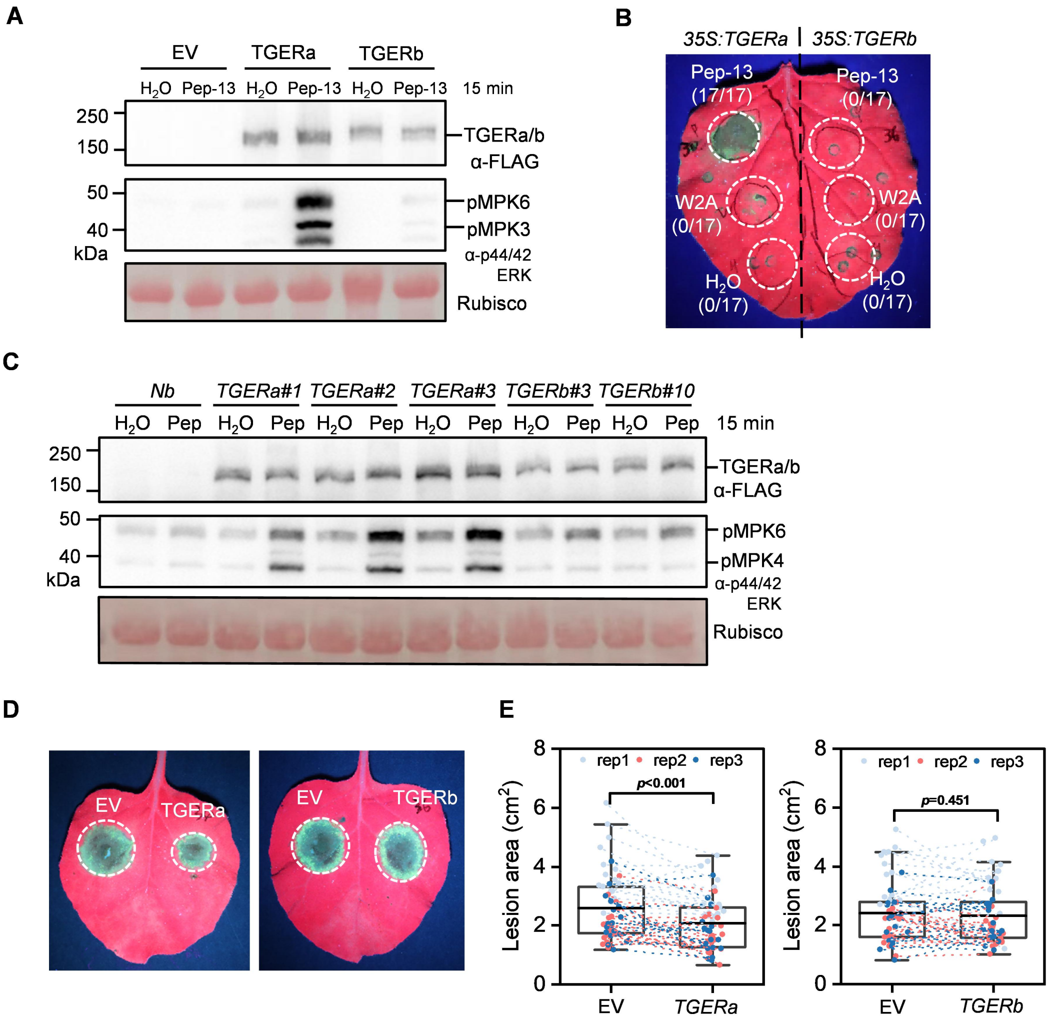


**Supplemental Figure 9. TGERa, but not TGERb, mediates MAPK activation and cell death in response to Pep-13, and confers enhanced resistance**

**(A-B)** Constructs harboring *35S:TGERa-3Flag* and *35S:TGERb-3Flag* were expressed in *Nb* leaves via *Agrobacterium*-mediated transient expression. After 24 h, 50 µM Pep-13 and $H_2O$ were infiltrated into leaves expressing *TGERa-3Flag* and *TGERb-3Flag*, respectively. MAPK samples were collected 15 min post-infiltration and analyzed by western blot using anti-Flag and anti-p44/42-ERK antibodies (A). For cell death analysis, 100 µM Pep-13, 100 µM Pep-13W2A and $H_2O$ were infiltrated as above. Cell death was observed only in *Nb* leaves expressing *TGERa-3Flag* at 24 h after Pep-13 treatment. Numbers in parentheses show the count of necrotic spots versus total spots (B).

**(C)** MAPK activation in *np-TGERa* lines (*TGERa #1*, *#2*, *#3*), *np-TGERb* lines (*TGERb #3*, *#10*) and *Nb* plants after treatment with 50 µM Pep-13 or $H_2O$.

**(D-E)** *TGERa* enhances resistance to *Phytophthora capsici* when transiently expressed in *Nb* leaves. (D)

Empty vector (EV), *35S-TGERa/b* were transiently expressed in *Nb* leaves for 24 h, followed by inoculation with an agar plug (5 mm diameter from colony margins of *P. capsici* strain BYA5. Detached leaves were incubated in the dark at 28°C for 2 days, and lesions were photographed under UV light. (D) Quantification of lesion areas using ImageJ. Data were analyzed using paired t-test (*TGERa*, $p < 0.05$, n=55 from three independent experiments; *TGERb*, $p = 0.451$, n = 56 from three independent experiments).

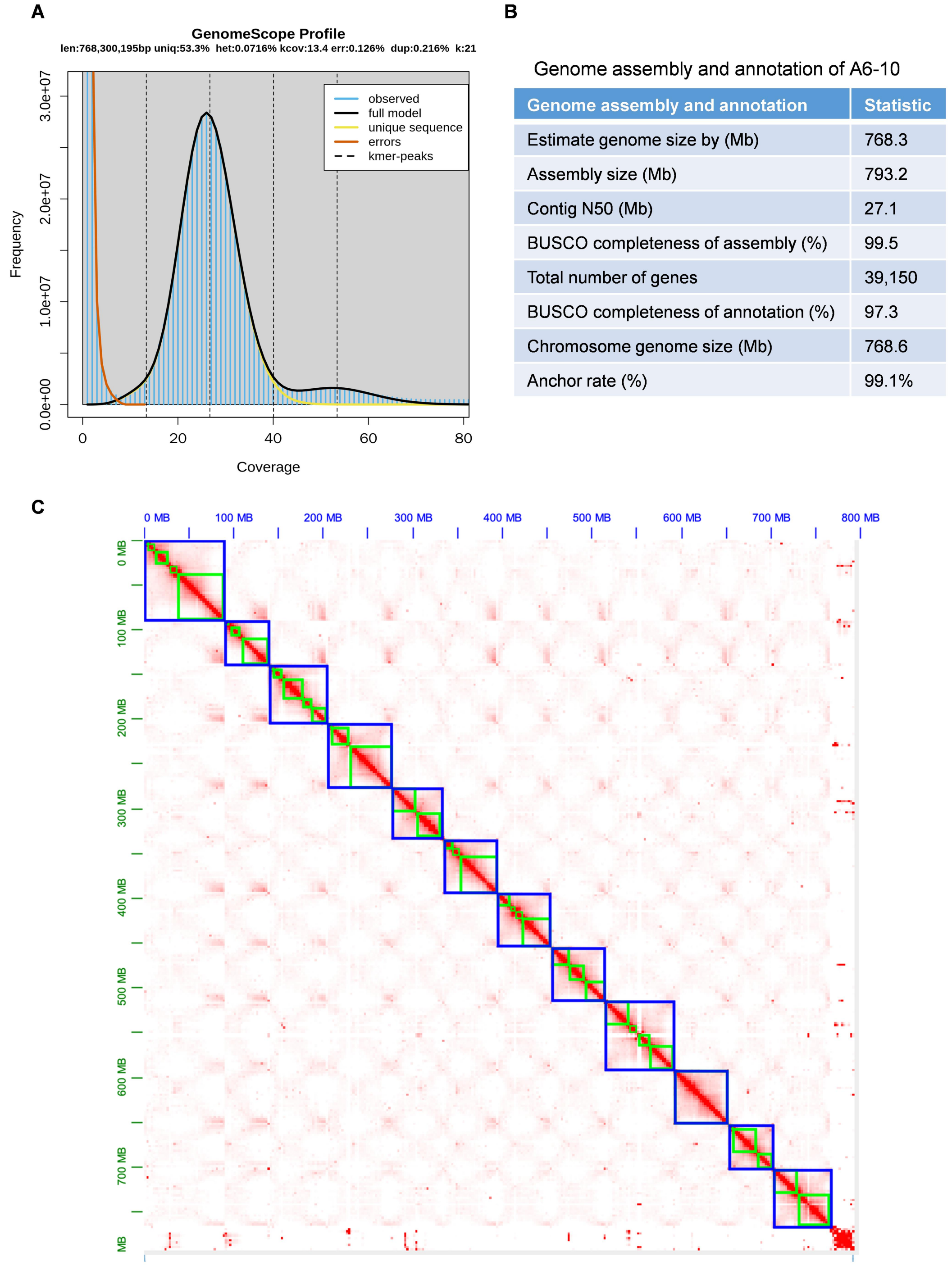


| Genome assembly and annotation | Statistic |
|---|---|
| Estimate genome size by (Mb) | 768.3 |
| Assembly size (Mb) | 793.2 |
| Contig N50 (Mb) | 27.1 |
| BUSCO completeness of assembly (%) | 99.5 |
| Total number of genes | 39,150 |
| BUSCO completeness of annotation (%) | 97.3 |
| Chromosome genome size (Mb) | 768.6 |
| Anchor rate (%) | 99.1% |

**Supplemental Figure 10. Genome assembly and annotation of potato inbred line A6-10.**

**(A)** K-mer spectrum analysis of A6-10 using HiFi sequencing reads and GenomeScope.

Abbreviations on the diagram: len – inferred total genome length, uniq – percent of the genome that is unique (not repetitive), het – overall rate of heterozygosity, kcov – mean k-mer coverage for heterozygous bases, err – error rate of the reads, dup – average rate of read duplications, k – k-mer size.
**(B)** Statistics summary of genome assembly and annotation for A6-10. The A6-10 genome data has been deposited at http://solomics.agis.org.cn/potato/species under Accession A6-10 (A6-10).
**(C)** Hi-C scaffolding for the A6-10 genome. Hi-C scaffolding heat map for the largest 12 chromosomes, showing chromatin interaction frequencies across the genome. Green blocks indicate contig positions, and blue lines denote chromosome boundaries, and color intensity represents the interaction frequency between different genomic regions.

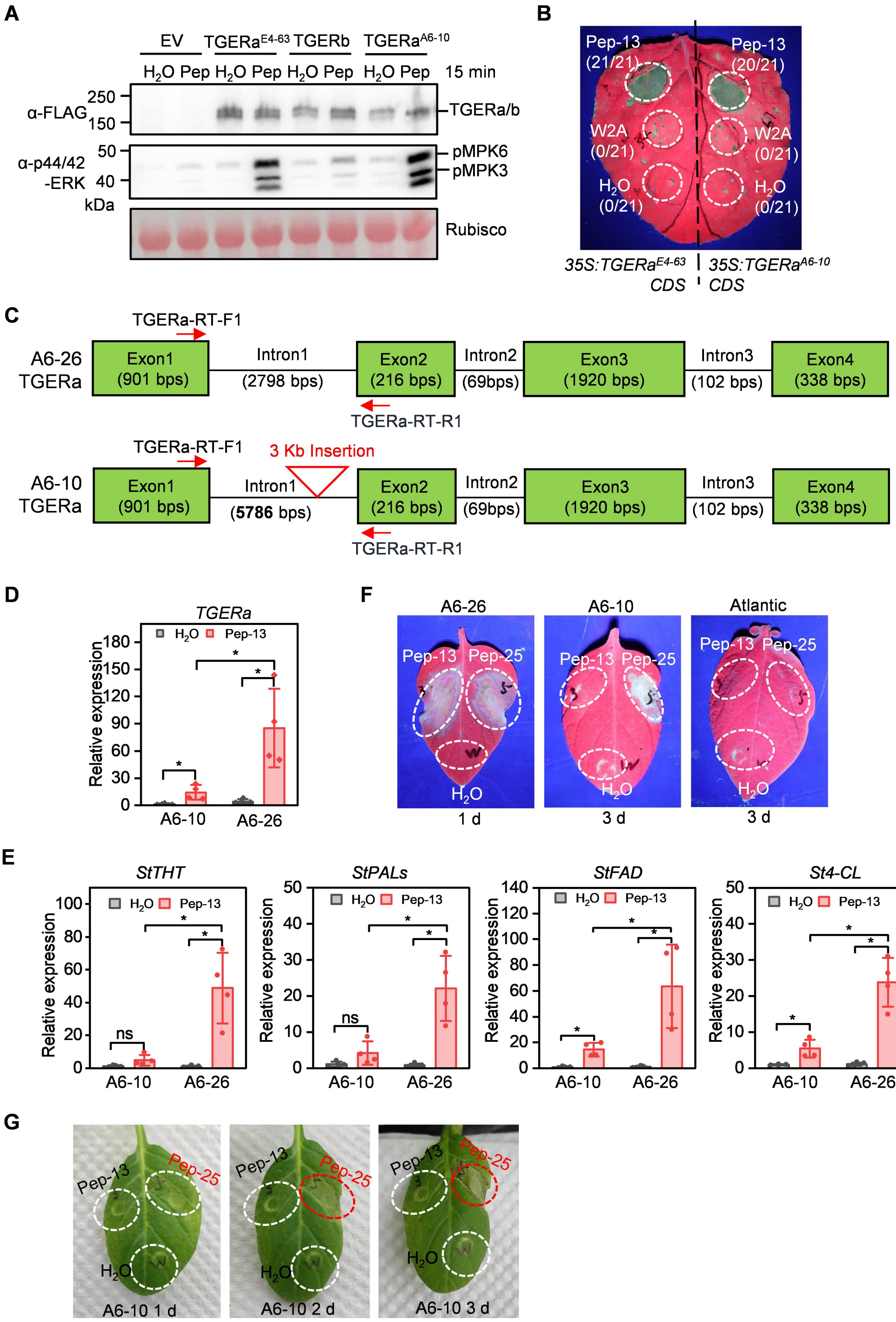


**Supplemental Figure 11. *TGERa* in A6-10 is a weak allele with reduced expression.**
**(A-B)** TGERa$^{A6\text{-}10}$ activate MAPK (A) and induce cell death (B) in response to Pep-13 when transiently expressed in *Nb* leaves. *TGERa*$^{E4\text{-}63}$ /*TGERb*/*TGERa*$^{A6\text{-}10}$-3Flag were transiently expressed in *Nb* leaves for 24 h. Leaves were infiltrated with 50 μM Pep-13 and $H_2O$. MAPK activation was analyzed in samples collected 15 min post-infiltration by western blot using anti-Flag and anti-p44/42-ERK antibodies. For cell death analysis, leaves expressing *TGERa*$^{E4\text{-}63}$*-3Flag* and

*TGERa*$^{A6\text{-}10}$*-3Flag* were infiltrated with 100 µM Pep-13, 100 µM Pep-13W2A and $H_2O$. Images were taken at 48 h post-infiltration.

**(C)** Gene structure of *TGERa* in the A6-26 and A6-10 genomes, with the sizes of exons and introns indicated. *TGERa*$^{A6\text{-}26}$ and *TGERa*$^{A6\text{-}10}$ share an identical exon–intron structure; however, a 3-kb DNA fragment insertion was identified in the first intron of *TGERa*$^{A6\text{-}10}$.

**(D)** *TGERa* transcript levels are strongly induced in A6-26 but show weak induction in A6-10. Transcript levels of *TGERa* were analyzed by RT-qPCR in leaves of A6-10 and A6-26 at 4 h after infiltration with 100 µM Pep-13 or $H_2O$. Data represent mean ± s.d. n=4, *$p$ <0.05, Student's t-test.

**(E)** Pep-13-responsive genes are strongly induced in A6-26 but show little or no induction in A6-10. Transcript levels of *StTHT*, *StPALs*, *StFAD*, and *St4-CL* were analyzed by RT-qPCR in leaves of A6-10 and A6-26 at 4 h after infiltration with 100 µM Pep-13 or $H_2O$. *NbEF1α* used as reference gene. Data represent mean ± s.d. n = 4. Asterisks indicate statistically significant differences (*$p$ < 0.05, Student's t-test); ns, no significant difference.

**(F)** Differential responses of A6-10, A6-26 and Atlantic to Pep-25. Potato leaves were infiltrated with 100 µM Pep-13, Pep-25, or water. A6-26 exhibited strong cell death as early as 1 day after infiltration with either Pep-25 or Pep-13. In A6-10, Pep-25 induced cell death at 3 days post-treatment, whereas Pep-13 failed to trigger cell death. Neither Pep-13 nor Pep-25 induced cell death in Atlantic.

**(G)** Time-course analysis of Pep-13- and Pep-25-induced cell death in A6-10. Leaves of A6-10 were infiltrated with 100 µM Pep-13, Pep-25, or water, and cell death phenotypes at the infiltrated sites were monitored over time. No visible cell death was detected in A6-10 at 1 day post-treatment. Cell death was triggered in Pep-25-infiltrated spots by day 2 and developed into distinct necrotic spots by day 3, whereas Pep-13 did not induce necrosis at any time point examined.
